# OmiXAI: An Ensemble XAI Pipeline for Interpretable Deep Learning in Omics Data

**DOI:** 10.1101/2025.04.28.651097

**Authors:** Ameliia Alaeva, Anna Lapteva, Natalya Mikhaylovskaya, Vladislav Malkov, Alan Herbert, Andrey Borevskiy, Maria Poptsova

## Abstract

Deep learning methods have become methods of choice in the analysis of genomic data. The performance of deep learning models depends on the information available for training. A growing trend in deep learning applications involves leveraging multi-omics data—spanning genomics, transcriptomics, epigenomics, proteomics, metabolomics, and other domains. When a deep learning model trained on omics data achieves high performance, the important question is to define factors that contribute to model’s predictive power. Explainable AI (XAI) methods can be categorized as model-aware and model-agnostic. Model-agnostic approaches, which rely on combinatorial feature perturbations to assess impact, are often computationally prohibitive for deep learning models. To address this, we developed OmiXAI, a pipeline integrating ensemble model-aware XAI methods. Our framework incorporates gradient-based techniques—including Integrated Gradients, InputXGradients, Guided Backpropagation, and Deconvolution (for CNNs and GNNs)—as well as Saliency Maps and GNNExplainer (specifically for GNNs). We evaluated OmiXAI on Z-DNA prediction using multi-omics features, demonstrating its efficacy through feature importance analysis and benchmarking of XAI methods. Notably, OmiXAI enabled feature engineering, reducing the critical feature set from almost 2,000 to just 50. Its modular design allows seamless integration of additional attribution methods, ensuring adaptability beyond omics to diverse problem domains. While testing the ensemble approach we benchmarked individual XAI methods and discuss their drawbacks and limitations. OmiXAI is freely available at https://github.com/aameliig/OmiXAI.

## Introduction

Deep learning methods proved to be efficient in predicting functional genomic elements such as promoters [1], enhancers [2], splice cites [3], flipons (non-B DNA structures) [4][5], histone marks [6][7], transcription factor binding sites [8], chromatin accessibilty regions [9][10], and others. Large Language Models in genomics achieved high performance by learning representations at the DNA sequence level and encorportaing contextual regions of different sizes [11][12][13][14][15][16]. Despite excellent results achieved by deep learning architectures trained on DNA sequence only, integration of omics data into machine learning algorithms not only improves a model’s prediction power but also helps to discover important associations between functional genomic elements and omics features [4][17][18]. Multi-omics approaches became particularly important in cancer research [19][20][21]. This brings the task of selecting the right explainable AI (XAI) methods to draw accurate conclusions of the highest importance.

Neural networks enable identification of complex non-linear relationships across a broad range of modalities. However, deep learning algorithms often operate as “black boxes” due to their intricate internal mechanics, comprising potentially millions of parameters with non-linear interactions. Understanding how models make their predictions and which relationships they identify as important can help elucidate the underlying mechanisms and processes. XAI methods have been progressing along with the development of deep learning algorithms and are still the subject of active research, especially in the field of bioinformatics, which usually adopts XAI techniques from other fields such as computer vision or natural language processing.

Interpretability of deep learning models in bioinformatics has been extensively reviewed [22][23][24]. In a systematic study of XAI methods [25] 48 papers (11.9%) were identified as using multi-omics data. however all the proposed methods are tailored for a specific task with a variable number and different types of omics data [18][26][27][28][29][30][31]. XOmiVAE is an interpretable deep learning model, based on variational autoencoder, for cancer classification using omics data (gene expression and CpG methylation profiles) [32]. DeepOmix is an interpretable multi-omics (DNA, RNA, proteins, metabolites) deep learning (CNNs, GNNs, autoencoders) framework with application in cancer survival analysis [33]. EUGENe [34] performs analyses of regulatory genomic sequences and uses several attribution approaches as interpretation methods. AutoXAI4Omics is model-agnostic but it is based on SHAP and Eli5 methods that are difficult to apply to deep learning models. See more methods reviewed in [25] and [35].

In this work we focus on model-aware XAI methods that can interpret how a well-trained deep learning model learned omics features space. Here we provide a concise overview of the fundamentals of state-of-the-art XAI methods that will be employed in this study.

The **Saliency** method, which is a common machine learning XAI method, employs gradients to identify input features that most significantly influence a network’s output [36]. The method leverages the fact that gradients reflect the sensitivity of the output to variations in the input. By visualizing these gradients, it becomes possible to highlight regions of an input image that have the greatest impact on the network’s predictions. Although several modifications have been proposed, the most common approach involves computing the gradient of the output with respect to the input image. This forms the foundation of gradient-based feature attribution methods in XAI. Despite its speed and simplicity, the Saliency method is prone to generating considerable noise. Subsequent modifications have sought to address this limitation by improving the method’s accuracy and reducing noise. Two subgroups of gradient-based, post-hoc feature attribution XAI methods to be described next do share similar but not identical relevance calculation strategies.

**InputXGradients** (IxG) introduces a minor modification to the original Saliency algorithm by proposing that the input data be multiplied by the calculated gradients [37]. This multiplication effectively amplifies the saliency of features that contribute most significantly to the network’s output, thereby making them more prominent for visualization.

The other two XAI methods - **Guided Backpropagation** (GB) [38] and **Deconvolution** (DC) [39] - also calculate the gradient of the output with respect to the input, but redefine the backward propagation of ReLU functions so that only non-negative gradients are propagated. In GB, the ReLU function is applied to the input gradients, whereas in DC, the ReLU function is applied to the output gradients, which are then directly propagated back. A key aspect of DC is the reverse application of operations such as convolution and pooling. Instead of simply computing the maximum during pooling, the method records the location of the maximum value during the forward pass and restores information only from these locations during the backward pass. This means that if a neuron was activated during the forward pass (i.e., its output was positive) and the gradient of this output with respect to the input is also positive, that gradient will be considered during the backward pass. If either condition is not met, the gradient is zeroed out. This approach allows for the identification of parts of the input tensor that directly contribute to the increased activation of the selected neuron, while ignoring those that decrease its activation.

The **Integrated Gradients** (IG) method builds on the Saliency method but enhances it by providing attribution scores that quantify the impact of each input feature on the model’s prediction [40]. Unlike to single-point gradient techniques, the IG method considers the influence of a feature along a continuous path in the input space. This path is constructed through linear interpolation between the original input and a designated baseline, often a zero-valued input representing a neutral state (e.g., a black image). The method involves calculating gradients at each point along this interpolation path. These gradients reflect the sensitivity of the model’s output to infinitesimal changes in each feature at a specific points along the path. The final attribution scores are obtained by integrating these gradients over the entire interpolation trajectory. This integration effectively averages the influence of each feature across the full range of values encountered during the interpolation process, resulting in more comprehensive and robust attribution scores compared to those generated by single-point gradient methods.

While the aforementioned approaches are applicable to both convolutional and graph neural networks, sharing similar propagation rules, **GNNExplainer** (GnnX) is specifically designed for graph neural networks [41]. The method identifies graph masks, which are effectively refined adjacency matrices that highlight the subgraphs and subsets of node features with the greatest influence on the model’s predictions. GNNExplainer achieves this by formulating an optimization problem that maximizes the mutual information between the distribution of GNN predictions and the distribution of possible subgraphs and subsets of node features.

A significant challenge in the XAI field is that different methods often yield inconsistent results. These discrepancies in feature importance rankings can be attributed to variations in the underlying algorithms and the unique characteristics of different neural network architectures. To tackle this, we present a novel feature importance pipeline OmixAI designed to identify key omics features that play a critical role in model predictions. The pipeline begins by individually training convolutional and graph neural network architectures. Although these architectures share sufficiently similar analytical strategies to allow simultaneous use and comparison, they also incorporate essential differences. Next, four gradient-based XAI methods — Deconvolution (DC), InputXGradients (IxG), Integrated Gradients (IG), and Guided Backpropagation (GB) — are applied to each trained network, and Saliency and GNNExplainer are applied to graph architecture. In the final stage, a hybrid ranking combines the feature importance generated by all XAI methods. For each feature, rankings are generated on the basis of statistical deviations from the mean, producing separate importance values for each model. The OmiXAI pipeline is designed to minimize the performance gap between the model’s initial accuracy and its performance after feature selection, ensuring efficient feature engineering. Additionally, it enables the identification of key feature dependencies within the studied biological context. We evaluate the effectiveness of OmiXAI pipeline on the Z-DNA identification with DeepZ model [17] based on omics features and compare the OmiXAI feature importance with those obtained with an enrichment method used in the study [17].

## Materials and Methods

### Z-DNA Data Set

We took the Kouzine dataset [42] assembled earlier in [4]. The method - permanganate/S1 nuclease footprinting [42] - can detect Z-DNA regions at base pair resolution and have more narrow regions compared to ChIP-seq peaks. The dataset consists of 45,201 Z-DNA intervals and includes 1,950 omics features such as transcription factor binding sites, histone marks, RNA polymerase binding sites, hDNase sites - all aggregated over different tissues, and dinucleotide energy transition from B- to Z-conformations (see full list in Supplementary Table 1).

### Omics Feature Selection and Preprocessing

We used the OmicsDC framework (https://github.com/hse-bioinflab/OmicsDC) that can automatically download and preprocess the omics features from ChIP-Atlas (http://chip-atlas.org) for a user-defined tissue and cell types. The DNA sequences are encoded using one-hot representations, omics features are integrated as additional features, and all the data is compressed using the SparseVector [4] module to optimize computational efficiency. Finally, all target labels are binary encoded, with a value of 1 indicating the presence of a corresponding functional element within the DNA regions.

### Deep Learning Model Training

Chromosome sequences were partitioned into fixed-length intervals, where interval size was treated as a tunable hyperparameter with the optimal size of 200 nucleotides. To ensure compatibility with existing deep learning methodologies for Z-DNA prediction [4][the dataset was constructed with a 3:1 ratio of negative (non-Z-DNA) to positive (Z-DNA-containing) intervals, where positive intervals contained at least some nucleotides classified as Z-DNA. To mitigate class imbalance, we employed stratified data partitioning based on both Z-DNA presence and chromosome identity.

All models were estimated with F-scores. To enhance interpretability, deep learning models should prioritize achieving a high rate of true positive (TP) predictions. Several deep learning architectures have been published for the task of Z-DNA prediction based on omics data[4][17][which were used as baselines for comparison.

Here, we designed CNN and GNN models that achieve the best performance. The CNN architecture, called ConvMZC (Figure 2A), consists of 12 convolutional layers (versus 3 in earlier DeepZ publications), group normalization, and alpha dropout. ConvMZC demonstrated a clear improvement in performance, with an F-score of 0.88 and an ROC-AUC of 0.97. Among the evaluated GNN architectures containing one of the renowned convolution techniques (GraphSAGEConv, GCNConv, GraphConv), the GraphMZC model (Figure 2B) contains 13 GraphSAGEConv layers, a neighbor sampling size of 400, and a class weight ratio of 1:25 for the loss function, and it achieved F-score of 0.81 and a ROC-AUC of 0.95 (Table 1).

**Fig. 1.**
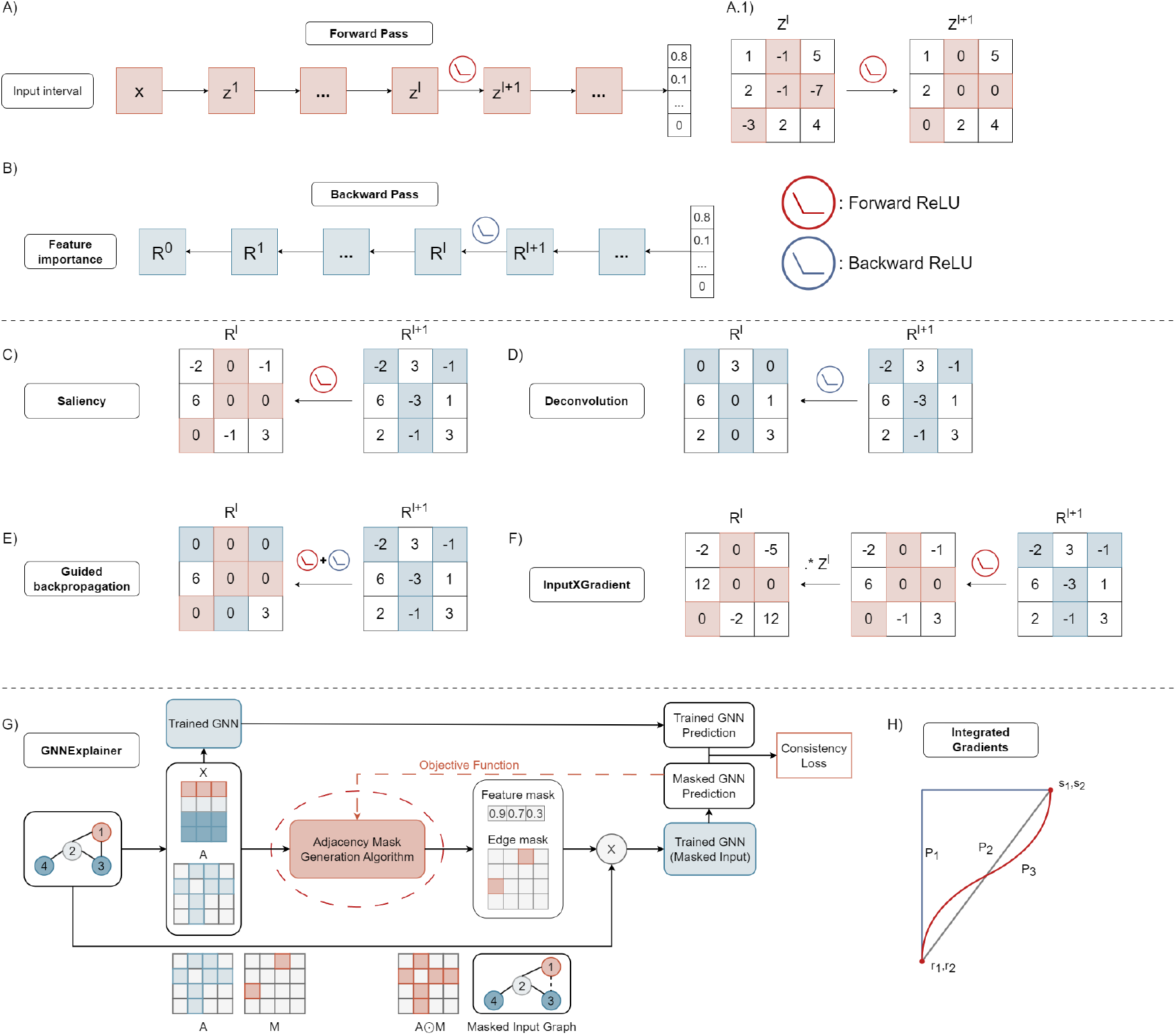
Scheme of feature attribution XAI methods. General methodology of (A) forward pass and (B) backward pass in neural networks; (C-F) Example matrices for the single forward and backward passes of XAI methods between *𝓁* and *𝓁* + 1 layers; (G) GNNExplainer (GnnX); (H) Integrated Gradients (IG)

**Fig. 2.**
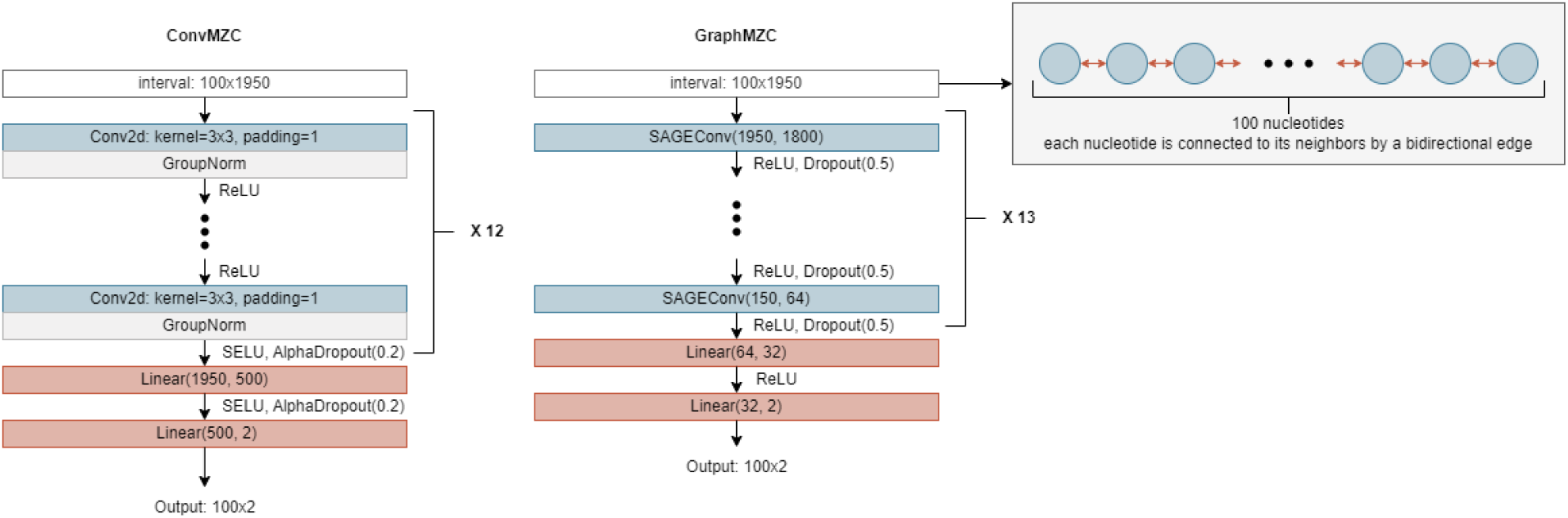
Architectures of best-performing convolutional and graph models for ZDNA classification task: (A) ConvMZC; (B) GraphMZC.

**Table 1.** Performance of deep learning architectures GraphMZC and ConvMZC.

| Model | Datset | Interval size | ROC-AUC | Precision | Recall | F1-score |
| --- | --- | --- | --- | --- | --- | --- |
| ConvMZC | Kouzine | 100 | 0.979 | 0.894 | 0.871 | 0.88 |
| GraphMZC | Kouzine | 100 | 0.958 | 0.796 | 0.829 | 0.81 |

### XAI Methods

Four gradient-based XAI methods - Integrated Gradients, InputXGradients, Deconvolution and Guided Backpropagation were applied separately to the trained convolutional and graph models. Additionally, two methods - Saliency and GNNExplainer were applied to only graph model. In total, the pipeline generated 10 feature importance values for each true positive (TP) region identified by two models, assigning feature-specific relevance scores to all 1,950 omics features. For XAI methods’ implementation we used the Captum library built for Pytorch [45]. Notably, OmiXAI is designed to be compatible with any XAI method, ensuring scalability and adaptivity.

Each of the four gradient-based methods was initialized with the target neural network provided as a single input parameter. Then the attribute method of the initialized object was called passing the model’s input tensor and target class as arguments. The model’s input tensor is a 2D tensor of shape (*N*_features_ *× N*_nucleotides_), and the target class is a tensor of size *N*_nucleotides_, comprising only ones. Attribute method returns the so-called input attributions, which have the same size and dimensionality as the inputs and contain the relevance scores for each of the features at each position across the entire interval. These relevance scores indicate how strongly each feature influenced the model’s decision toward the positive class at every nucleotide position within the interval.

For GraphMZC, we additionally utilize the explanation module from PyTorch Geometric, which provides the specialized GNNExplainer algorithm for graph models and enables the application of Captum methods to graph architectures. We recommend using the four gradient-based methods mentioned above, along with Saliency and GNNExplainer, for graph neural network interpretation. The interpretation pipeline remains consistent, with the only difference being the initialization of a GNNExplainer object configured with the target model, the selected interpretation algorithm, and the algorithm’s default settings as properties.

## Results

### Feature selection pipeline

Here we present feature selection pipeline based on model-aware XAI methods that are applicable to deep learning models that incorporated omics data in network architectures.

The framework consists of the following blocks: Data Preprocessing, Model Training, Interpretation, and Hybrid Ranking. The general schema of the pipeline is presented in Figure 3.

**Fig. 3.**
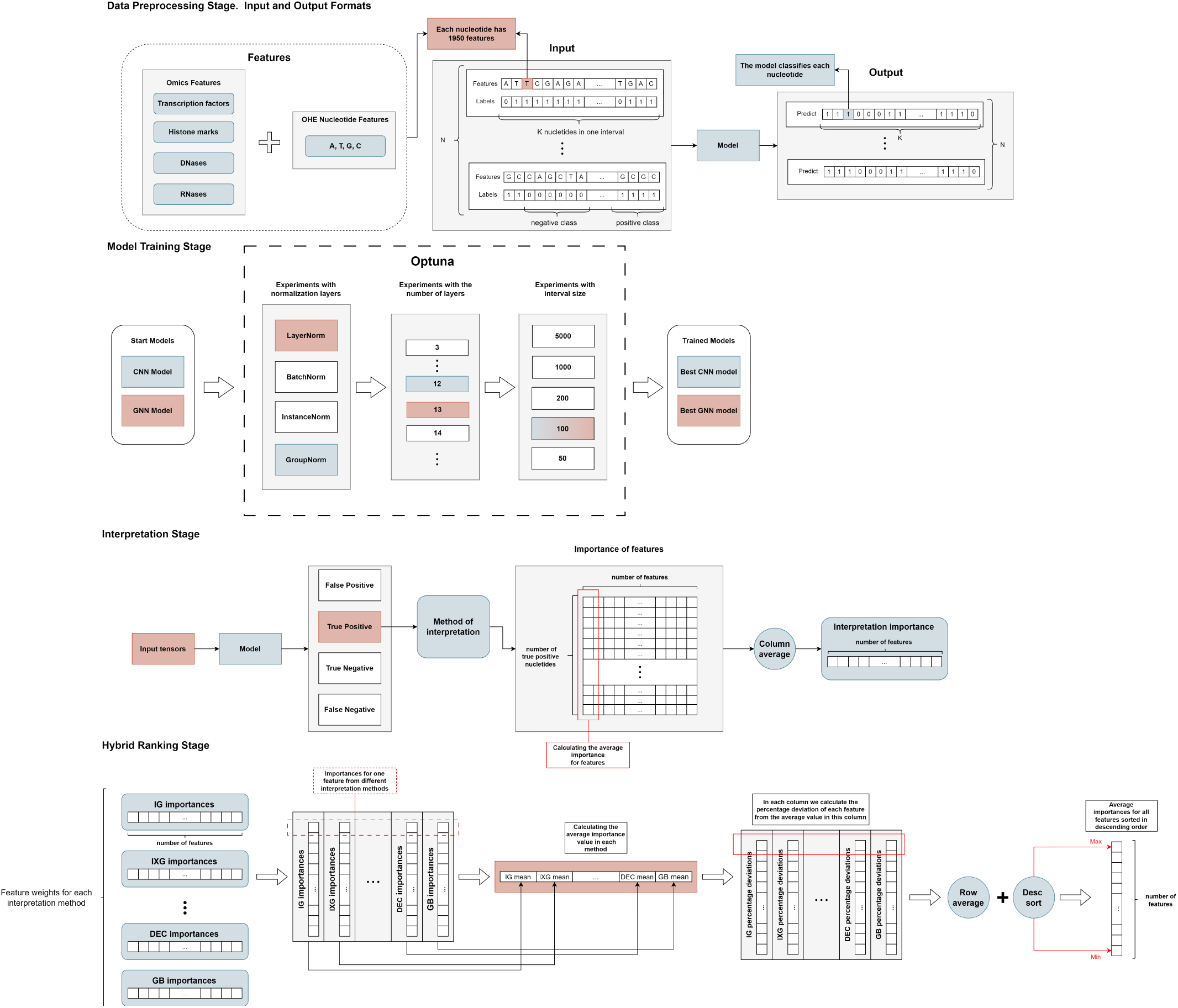
Feature Selection pipeline scheme: (A) Data Preprocessing Stage (B) Model Training Stage (C) Interpretation Stage (D) Hybrid Ranking Stage

### Data Preprocessing

Functional genomic elements are presented as a DNA sequence with corresponding omics features that can include histone marks, transcription factor binding sites, chromatin states, Hi-C maps, methylation level, and any other omics experiments (see Methods for omics feature preprocessing).

### Model Training

At this stage, two complementary neural networks - convolutional and graph architectures - are developed. Each architecture is independently designed to solve the classification task using nucleotide data and corresponding omics features. Particular attention is paid to identifying regions associated with the positive class, as these regions play crucial role in subsequent interpretation and analysis.

### Interpretation

A post-hoc interpretation step is implemented to identify and analyze the features that significantly contributed to the model’s accurate prediction of true positives (TPs). This is achieved using two analogous groups of model-aware gradient-based interpretation methods. Here we use Integrated Gradients, InputXGradients, Deconvolution, Guided Backpropagation, Saliency and GNNExplainer. All methods finally produce feature importance values that are combined together at the next stage.

Interpretation is similar across all selected XAI methods. First, a single object is processed to make a prediction, identifying the indices corresponding to the TP region. Next the unchanged input tensor is passed through an interpretation algorithm, focusing on the target class. The indices previously identified as TPs are preserved throughout this process.

### Hybrid Ranking

This stage focuses on prioritizing features based on the averaged interpretation results from each XAI method. At the end of the previous step, the pipeline generates ten importance scores for each of the features. The next step is to rank the features separately for each model, combining the statistics gathered from different XAI approaches. The final ranking is calculated as follows:

1. Compute the average of all the relevance scores of each XAI method.
2. For each object, determine the percentage deviation of its relevance score from the corresponding mean for each XAI method. All relevance scores for each feature are thus transformed into the same number of percentage deviations.
3. Calculate the mean percentage deviation of each feature across all XAI methods for the given model. As a result, each feature will have two average values: one from CNN and one from GNN.
4. Sort the list of mean percentage deviations in descending order, generating two independent rankings - one for CNN and one for GNN. This enables a comparison of the similarity between the two rankings and an assessment of the relative importance of each omics feature across the two distinct yet related architectures.

The interpretation pipeline produces the ranking list of feature importance.

### Application of OmiXAI to Z-DNA

#### Omics Feature Importance Analysis with OmiXAI

The developed interpretation pipeline can be used for any functional genomic elements. In this study, we demonstrate its performance for Z-DNA regions, as deep learning models trained on omics data for Z-DNA prediction are currently available and published together with the omics importance analysis performed by other methods [4][17].

Using Z-DNA dataset with 1,950 features we trained ConvMZC and GraphMZC (see Methods for details). At this stage, only the intervals containing functional genomic elements of interest (here Z-DNA regions), i.e. true positives are retained. Each TP Z-DNA region are processed with XAI methods, each generating specific relevance score for each of the 1,950 features associated with the Z-DNA region. By repeating ConvMZC and GraphMZC inference for each Z-DNA region, we obtain 10 importance scores for each feature: four with ConvMZC and six with GraphMZC.

In total 45,201 tensors were interpreted. The set of acquired values along each XAI method enables the calculation of an average value, thus providing each feature with a specific mean importance score (Supplementary Table 1).

Correlation of top-1000 omics features ranked by each XAI method for GNN and CNN models is presented in Figure 4. We see that the highest similarity is observed between XAI methods applied to GNN as compared to CNN. The lowest intersection in feature ranking is for Deconvolution and Guided Backpropagation applied to CNN and between all the other methods. In general inside GNN group correlation between different XAI methods is higher than inside CNN group.

**Fig. 4.**
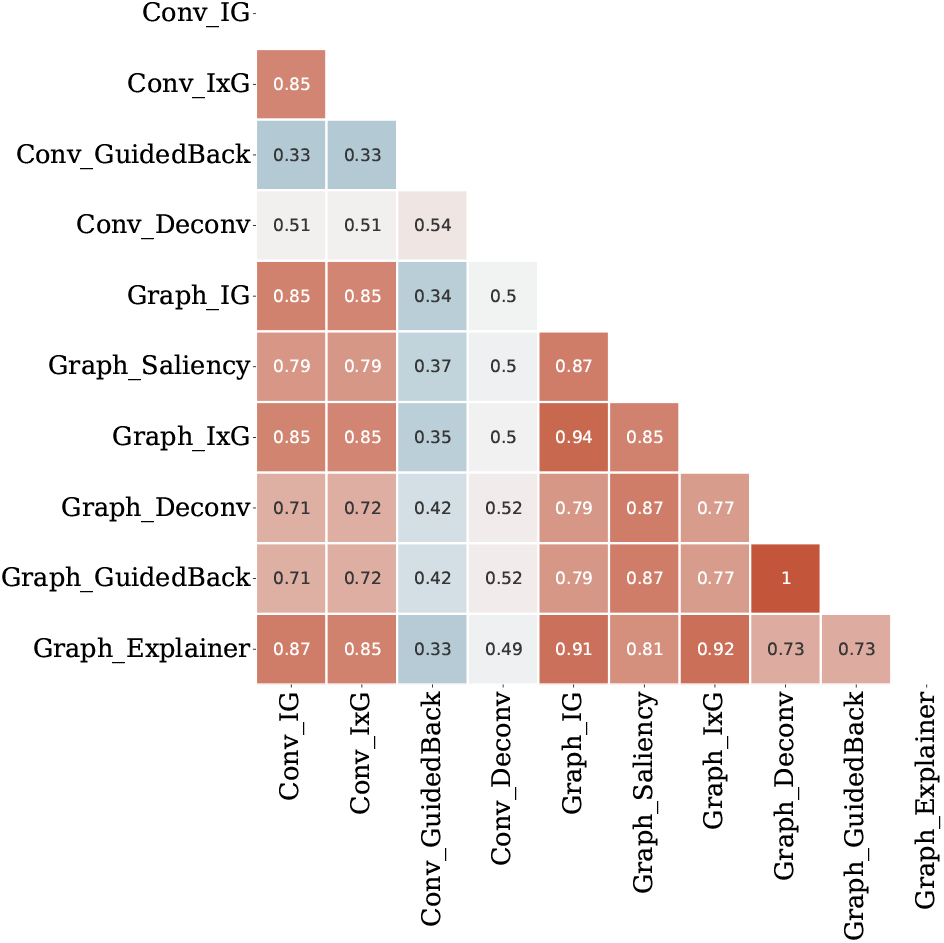
Correlation between feature importance generated by XAI methods for GraphMZC and ConvMZC. Methods for CNN are designated with Conv and for GNN are designated with Graph.

We applied Hybrid Ranking to the table with relevance scores and the final feature importance results generated by the pipeline are presented in Supplementary Table 1. Top 20 features include sites of RNA polymerase II; transcription factors CTCF, ERG, MAX, BCOR, TRIM25, MYCN, SUZ12; histone marks H2A.Z, H3K4me1,H3K4me2, H3K27ac; methylation 5-mC (Figure 5A) - all inline with the previous studies [4][17].

**Fig. 5.**
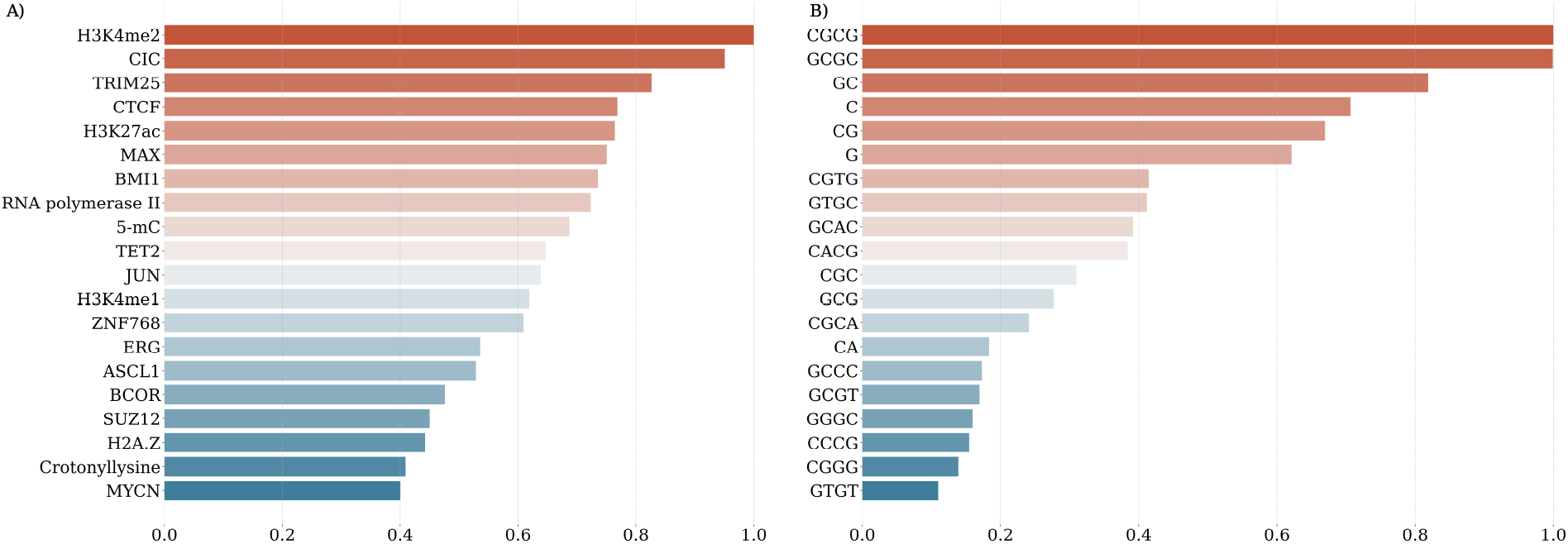
OmiXAI Feature Importance Plot for Z-DNA detection based on omics features (A) and k-mers as features (B).

#### Validation of Important Features

To evaluate how feature reduction affects model performance, we trained ConvMZC and GraphMZC on subsets of omics features ranked by importance (Table 2 and Table 3). The results demonstrate that the number of omics features can be reduced to 50 without losing predictive power. Interestingly, both models exhibit a slight increase in F-score when using 300 features, suggesting an optimal balance between dimensionality and information retention. This implies that when analyzing aggregated omics data (as opposed to tissue-specific datasets), only a small subset of omics features remains critical—most can be discarded while preserving key features alongside nucleotide-level information.

**Table 2.** Performance of retrained ConvMZC architecture on Kouzine dataset with top-*k* features and interval size of 100 nucleotides.

| *k*, № of top features | ROC-AUC | F1-score |
| --- | --- | --- |
| 1950 | 0.9789 | 0.88 |
| 704 | 0.9755 | 0.879 |
| 504 | 0.9778 | 0.88 |
| **304** | **0.9771** | **0.8815** |
| 104 | 0.9739 | 0.8668 |
| 54 | 0.9748 | 0.8682 |

**Table 3.**
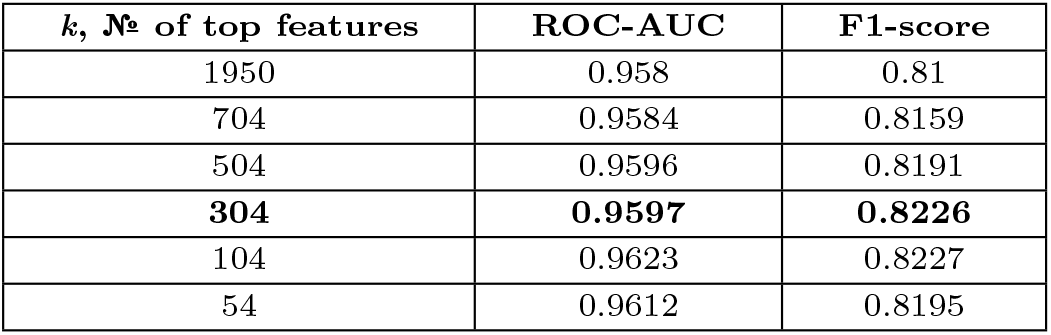
Performance of retrained GraphMZC architecture on Kouzine dataset with top-*k* features and interval size of 100 nucleotides.

| *k*, № of top features | ROC-AUC | F1-score |
| --- | --- | --- |
| 1950 | 0.958 | 0.81 |
| 704 | 0.9584 | 0.8159 |
| 504 | 0.9596 | 0.8191 |
| **304** | **0.9597** | **0.8226** |
| 104 | 0.9623 | 0.8227 |
| 54 | 0.9612 | 0.8195 |

**Table 4.** Performance of retrained GraphMZC architecture on Kouzine dataset with k-mers features and interval size of 100 nucleotides.

| $k$, № of features | k-mer | F1-score |
| --- | --- | --- |
| 20 | 1-2 | 0.856 |
| 84 | 1-3 | 0.857 |
| 340 | 1-4 | 0.866 |
| 1364 | 1-5 | 0.858 |

#### Omics Feature Analysis

In the work of [17] omics feature importance analysis was done with permutation feature importance (PFI) method for Z-DNA conserved between human and mouse and based on the set of common omics features. We intersected top-300 features detected by hybrid rankings for convolution and graph models with top-300 PFI important features and obtained 185 (61%) common features. It should be noted that the total number of omics features used in DeepZ model in [17] was 537 that is common for human and mouse, and the model evaluated here used 1,950 features so that the comparison is not straightforward. And permutation feature importance analysis also has its bias in the number of available samples for a particular omics feature, tissue representability, and other statistical factors. Here we think that OmiXAI pipeline is more reliable in feature importance ranking than statistical permutation tests for enrichment.

#### OmiXAI for k-mers importance analysis

Here we applied OmiXAI pipeline to k-mer importance analysis by replacing omics features with k-mers. For that we re-trained convolutional and graph neural networks with the novel input data of k-mers and applied hybrid ranking as described in the Methods. As a result of the OmiXAI feature importance analysis, the top-20 most important k-mers include CGCG, GCGC, GC, CG, CGTG, GTGC, GCAC, CACG, CGC, GCG, CGCA, CA, GCCC, GCGT, GGGC, CCCG, CGGG, GTGT (Figure 5B), which are indicative of Z-DNA formation (see full ranking in Supplementary Table 1). The results of this test provided further confirmation that the methodology is valid and adaptable to other functional genomic elements.

## Discussion

Deep learning models consistently outperform traditional tree-based approaches in predicting functional genomic elements from multi-omics data. This superior predictive performance, however, comes at the cost of interpretability, creating a critical need for reliable XAI methods specifically designed for deep learning architectures in genomic applications. Model-independent methods like SHAP or LIME are difficult to apply to deep learning models because they are based on perturbation of features and are computationally intensive. The number of possible feature combinations grows exponentially with the number of features and SHAP or LIME are impractical to apply to deep learning models. In this case, go inside architecture seems more advantageous.

For many genomic tasks, nucleotide or a region of nucleotide can be represented as a vector in a space of omics features and for those tasks gradient-based XAI methods seems to be the methods of choice. Gradients capture how small changes in input affect the output, making them useful for identifying influential features. Since gradients are already computed during backpropagation, gradient-based attribution methods are more fast and scalable compared to perturbation-based approaches (e.g., LIME or SHAP).

All of the presented interpretation methods focus on determining which input features have the greatest influence on the network’s output, but they interpret importance differently depending on how they handle gradients. Saliency simply takes the gradient of the output with respect to the input, so it can be seen as a first-order approximation of how each feature affects the output. Deconvolution also computes the gradient with respect to the input but treats the ReLU activation differently by propagating only positive error signals backward. Guided Backpropagation is similar to Deconvolution but goes a step further, combining vanilla backpropagation with the principle of “only positive gradient,” meaning it zeros out both negative gradients and negative inputs. InputXGradient extends the Saliency approach by multiplying the raw gradients by the actual input values. This is particularly intuitive for linear models, where the gradient corresponds to each feature’s coefficient, and “an input multiplied by a coefficient” provides insight into each feature’s overall contribution. Although these methods share the same goal — to show how each input feature affects the model’s predictions — the specific ways in which they backpropagate and combine gradients can lead to different representations and interpretations. Ensembling these methods can mitigate outliers produced by individual approaches while consensus features are reinforced.

Neural network interpretation algorithms have inherent drawbacks that limit the reliability of the obtained relevances unless hyperparameters and input data are precisely tuned. Gradient-based methods depend on the choice of baseline, can produce erroneous results if the network output saturates in the early stages of integration, and often involve high computational costs-especially for Integrated Gradients [46] [47]. InputXGradients inherits the problem of gradient saturation (for example, when using ReLU) and can be very sensitive to noise, highlighting features with negligible relevance [48] [49]. Methods such as Guided backpropagation and Deconvolution share a common drawback: they can produce visually appealing “edge detection” saliency maps that remain almost unchanged even if the network weights are randomized, indicating that their results often reflect general input features rather than representations learned by the model [50]. In more complex interpretation tasks, these algorithms can miss critical object details or incorrectly distribute relevances, raising questions about their reliability [51] [52] [53]. More generally, gradient-based interpretation tends to be unstable with minor changes to the input data [54].

In this approach, we can assess the extent of this “clever Hans” problem by using experimentally validated data to demonstrate this issue [55]. For example, the top “omic feature” is listed as DDX5 association with Z-DNA. However, genome-wide assays reveal this interaction is enriched over the Short Interspersed Repeat Element (SINE) Alu repeat. There are over one million SINEs copies in the human genome [56]. Due to mapping problems, Kouzine et al excluded them from the dataset used to identify Z-flipons in the genome, so it would not be expected that proteins mapping to SINEs would appear so prominently in the results. Similarly, CG islands are overrepresented in the genome in promoter regions and contain both Z-flipons and G-flipons. The high ranking of NPM1, a validated G-quadruplex binding protein [57], reflects this misattribution problem. Finding a strategy to overcome this critical limitation of xAI is essential to further developing these methods. For now, results should be interpreted cautiously. The findings require careful validation against existing experimental datasets.

Thus each of the benchmarked method can produce erroneously high importance for a particular feature, requiring additional caution when drawing scientific conclusions. However our experiments with reduced number of features ranked by importance showed that XAI methods can effectively reduce features space by eliminating less important.

The developed OmiXAI pipeline demonstrated performance for CNN and GNN however it will also work for transformer architecture. One can compute gradients of the output with respect to the input tokens/features, and one can also compute gradients with respect to attention weights, which gives insights into which parts of the sequence the model is “looking at” to make predictions.

In omics data research, where accurate interpretation of sequences can lead to significant biological discoveries, the use of an ensemble of methods helps cross-verify important features, reduce the influence of method-specific artifacts, and ensure that the discovered relevances genuinely affect the network’s predictions.

## Conclusion

Gradient-based xAI methods are efficient, precise, and model-aware, making them a popular choice for interpreting deep learning models while balancing computational cost and interpretability. We propose an integrated approach to extract key features from the complete feature map, revealing biological relationships relevant to specific genomic investigations. These extracted features, primarily omics data, are critical for deep neural network predictions, enabling the discovery of previously undocumented relationships. Additionally, OmiXAI pipeline not only identifies the most significant features but also ranks them, providing a feature engineering technique to reduce the feature set size required for training neural networks without compromising model quality.

OmiXAI is broadly applicable, extending beyond omics to various problem domains. By leveraging the complementary architectures of convolutional and graph neural networks alongside gradient-based interpretation methods, it aims to identify efficient pathways for selecting the most impactful features for target class identification.

## Data availability

OmiXAI is freely available at https://github.com/aameliig/OmiXAI.

## Author contributions statement Acknowledgments

